# Macroecology of microbial performance across Earth’s biomes

**DOI:** 10.64898/2026.01.30.702831

**Authors:** Angel Rain-Franco, Adrian-Stefan Andrei, Jakob Pernthaler

## Abstract

Microorganisms colonize all environments on Earth, yet it remains unclear which taxa merely broadly persist and which consistently outperform others across environments. Here we introduce the Baas-Becking score (BB-score), a performance metric to rank taxa across communities that integrates occupancy, relative abundance, and penalized absences. Applying BB-scores to 576,531 microbial communities spanning 24 biomes (categorized as either host-associated or free-living), we found that most species-level operational taxonomic units (OTUs) were widespread, but their success was substantially more restricted. Although 64% of OTUs were detected in at least one host-associated and one free-living biome, only six taxa ranked within the top 1% of performers in >50% of biomes: *Aerococcus viridans, Faucicola* (previously: *Moraxella*) *osloensis, Lawsonella clevelandensis, Methylorubrum populi, Sphingobium yanoikuyae,* and *Pseudomonas fluorescens* complex. These globally successful taxa were present in a quarter of all airborne communities, consistent with the atmosphere acting as a dispersal corridor. Network analysis of shared top 5% performers identified the phyllosphere and freshwaters as hubs linking animal-associated, plant-associated, and soil biomes. BB-score provides a scalable framework to map microbial success across Earth’s biomes and to put new focus on globally successful yet woefully understudied taxa.

**Signficance:** Microbes shape ecosystems, human health, and global biogeochemical cycles, yet we still lack tools to distinguish ecological success from occupancy. Here, we introduce a framework to rank taxa across communities, the Baas-Becking score (BB-score). Applying this framework to more than half a million microbial communities spanning 24 host-associated and free-living biomes, we show that strict biome specialization is uncommon and that occupancy rarely translates into high performance. Our results further reveal that Earth’s systems are connected through their microbiomes, with the atmosphere, plant surfaces, and freshwater systems representing key hubs. This perspective shifts microbial macroecology from mapping distributions to understanding ecological success and connectivity.

## Introduction

Microorganisms drive biogeochemical cycling in every environment on Earth (1). They sustain key ecosystem functions that regulate soil fertility, water quality, and climate, with cascading effects on agricultural productivity and plant and animal health (2, 3). Standardised, global sequencing initiatives such as the Earth Microbiome Project (4) have laid the foundation for macroecology studies by revealing large-scale spatial structure in microbial diversity and community composition. In such analyses, biomes often serve as the primary organizing framework for environmental comparisons. Originally defined as major climate zones (5), biomes are increasingly treated as higher-level ecological units that group communities shaped by similar environmental filters and shared habitat architecture and connectivity (vegetation structure, soil matrix, hydrologic networks) (6). This spatial framework has revealed consistent macroecological patterns, including rank-abundance microbial taxa distributions and systematic differences in alpha, beta, and gamma diversity patterns between aquatic and terrestrial systems (7, 8). Yet, while such community-level summaries quantify richness, diversity, and turnover, they do not capture how individual microbial taxa perform relative to others across major environmental discontinuities.

Despite immense microbial diversity, a small subset of taxa dominates within many biomes: approximately 2% of soil phylotypes account for about half of total abundance (9), SAR11 and a few cyanobacterial lineages dominate ocean planktonic biomass (10, 11), and relatively few taxa prevail across host- and plant-associated habitats such as the gut, skin, phyllosphere, and rhizosphere (12) (13) (14). However, these “winners” are largely biome-specific, motivating a unified, quantitative framework to quantify taxon success across environments and identify top cross-biome performers.

Attempts to assess the performance of taxa typically dichotomize into “abundant” and “rare” microorganisms; while the former often drive key ecosystem processes, the latter contribute to compositional and functional diversity (15, 16). Few studies have combined both metrics (abundance and occurrence) into a coherent measure (17). Here, we develop and implement an index that integrates both occurrence and relative abundance changes of individual taxa across communities. It is based on the Elo-rating system, originally designed to rank chess players (18). Elo ratings update across multiple communities, yielding a rank-based score for taxon changes, thereby providing a comparative measure of ecological success (19). We changed one assumption of the original Elo framework (“you don’t play, you don’t lose”) by incorporating taxa absences (“you don’t play, you lose”) into a non-zero-sum ranking parameter, thereby obtaining a scalable metacommunity phylotype-level index of relative performance. In recognition of L. Baas Becking’s influential proposition that “everything is everywhere, but, the environment selects” (20), the modified Elo rating parameter was designated the BB-score. The BB-score advances microbial macroecology beyond conventional diversity summaries by ranking taxa against their local competitors across communities, enabling direct performance comparisons across biomes, disentangling ubiquity from dominance, and identifying hub environments that promote cross-biome connectivity.

## Results

We queried the Microbe Atlas Project (MAP) database (21) and classified samples into 24 biomes of host-associated and free-living lifestyles, using primary and secondary habitat descriptors (Suppl. Table S1). The resulting dataset comprised 576,531 microbial communities with global coverage across terrestrial, aquatic, and host-associated environments (Fig. 1A). Animal-associated (including human) microbiomes accounted for 72.3% of samples, followed by freshwater (11.9%), soils (7.1%), plant-associated (4.3%), saline (4.1%), and airborne communities (0.3%; Fig. 1B, Suppl. Table S2). The full dataset comprised 908,147 bacterial and archaeal operational taxonomic units (OTUs) clustered at 99% sequence identity (22). Although sampling effort was concentrated in animal-associated biomes, the distribution of OTU richness across biome groups was markedly less skewed: animal-associated biomes contributed 23.4% of total OTUs, freshwater sytems 26.5%, soils 21.2%, plant associated 14.0%, saline systems 12.0%, and airborne 2.7% (Fig. 1C).

**Figure 1.**
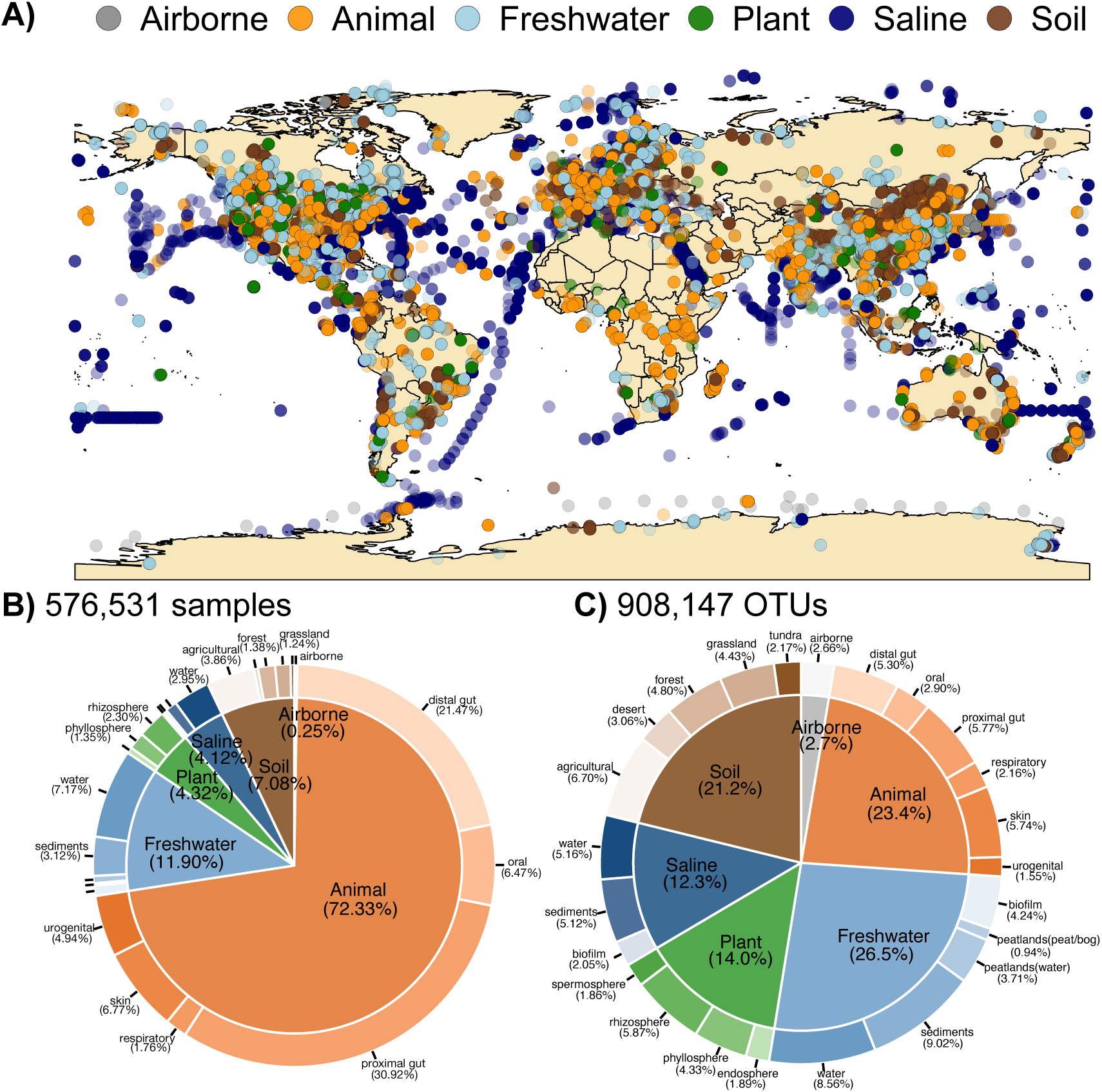
Sample set and biome classification for microbial performance assessment. (A) Global distribution of microbial community samples from animal, plant, soil, and water biomes derived from the Microbe Atlas Project (MAP dataset). (B) Distribution of samples among biomes. (C) Distribution of 16S rRNA gene based operational taxonomic units (OTUs, 99% identity) among biomes. In B-C, inner rings show main biomes classification (n=6), while outer rings show sub-biome categories (n=24).

### Global microbial occupancy

To quantify OTU occupancy across biomes while controlling for unequal sampling effort (Fig. 1B), we standardized each biome to the smallest one (plant spermosphere; n = 1,059 samples). We randomly drew 1,059 samples per biome without replacement and repeated this procedure 1,000 times (Methods). For each OTU, we quantified the number of distinct biomes in which it was detected in each subsampled dataset and summarized occupancy across iterations.

Across the 24 biomes, we identified 124,772 unique OTUs, most of which spanned both fundamental lifestyle categories: 63.8% were detected in at least one host-associated biome and at least one free-living biome, whereas 24.9% and 11.3% were restricted to free-living or host-associated biomes, respectively (Fig. 2A). Occupancy was strongly right-skewed. Only 2.58% of OTUs (3,223) were detected in more than 20 biomes, whereas 11.2% were consistently confined to a single biome across all subsampling iterations (“single-biome OTUs”), most prominently in freshwater and saline systems and in animal guts (Fig. 2A). Single-biome OTUs comprised only a small fraction of local diversity (median 0.92% of each biome’s OTU pool; IQR 0.47– 1.43%), ranging from ∼4% in freshwater sediments to <0.3% in airborne, peatland, and spermosphere biomes (Fig. 2B).

**Figure 2.**
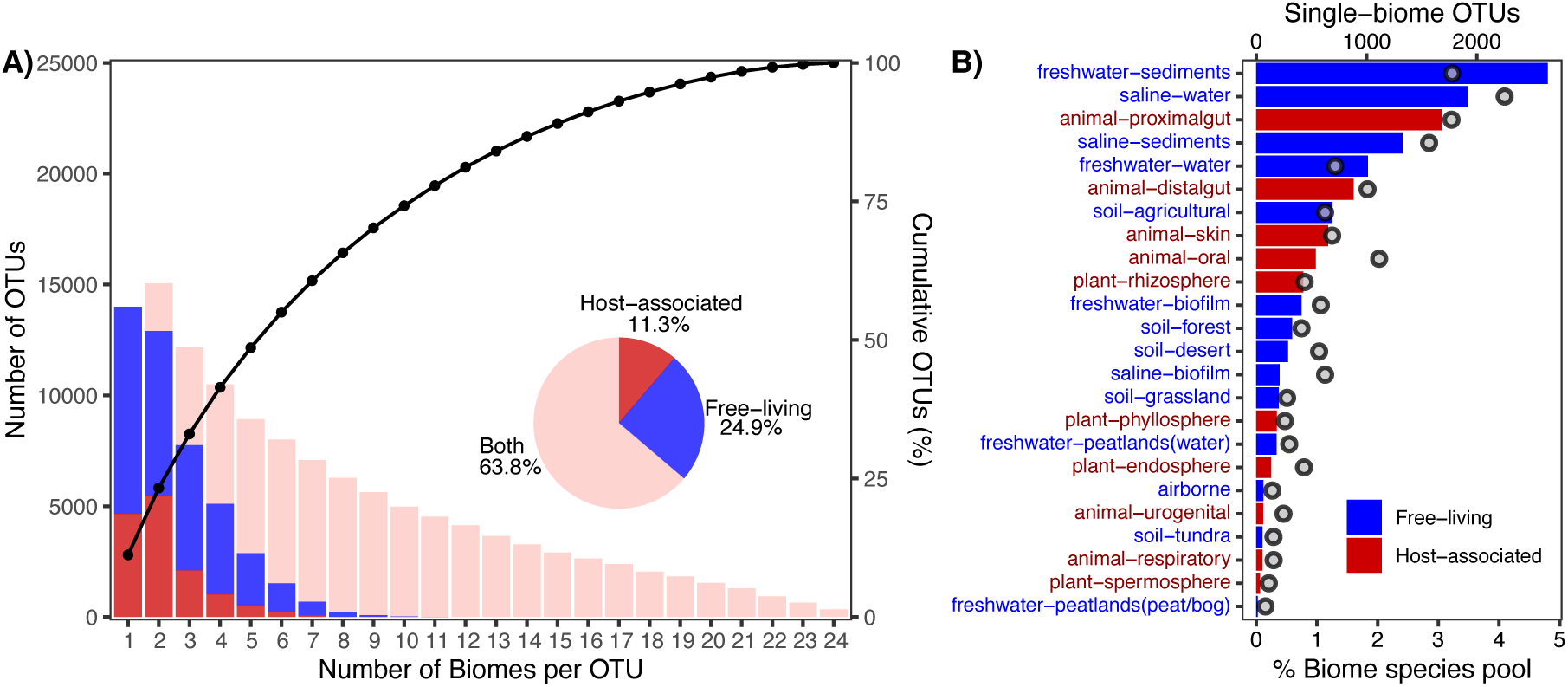
Occurrence of OTUs in biomes according to lifestyle. (A) Distribution of OTUs according to number of biomes in which they were found. Bars (left axis): number of OTUs occurring in one or more biomes (see Methods). Line and symbols (right axis): cumulative percentage of OTUs in increasing numbers of biomes. Inset pie chart: proportions of OTUs exclusively present in host-associated or free-living biomes, or found in both types. (B) Contribution of host-associated and free-living OTUs that are limited to a single biome to the total biome species pool. Bars (top axis): number of OTUs; grey symbols (bottom axis): proportion of total species pool of the respective biome.

### Assessing microbial performance by the BB-score

The BB-score distributions of OTUs within biomes were always right-skewed, indicative of a small number of high-performing lineages against a large majority of modest or low performers (Fig. 3A). Free-living biomes generally had higher mean BB-scores than host-associated ones (Wilcoxon test, W = 127, P = 0.002). Mean biome BB-scores were positively correlated with various alpha-diversity metrics such as species richness (Fig. 3B), Simpson index (Fig. 3C), and Pielou’s evenness (Fig. 3D). There was no significant relationship of BB-scores with OTU numbers per biome (Spearman ρ = 0.363, P = 0.082; Fig. 3E).

**Figure 3.**
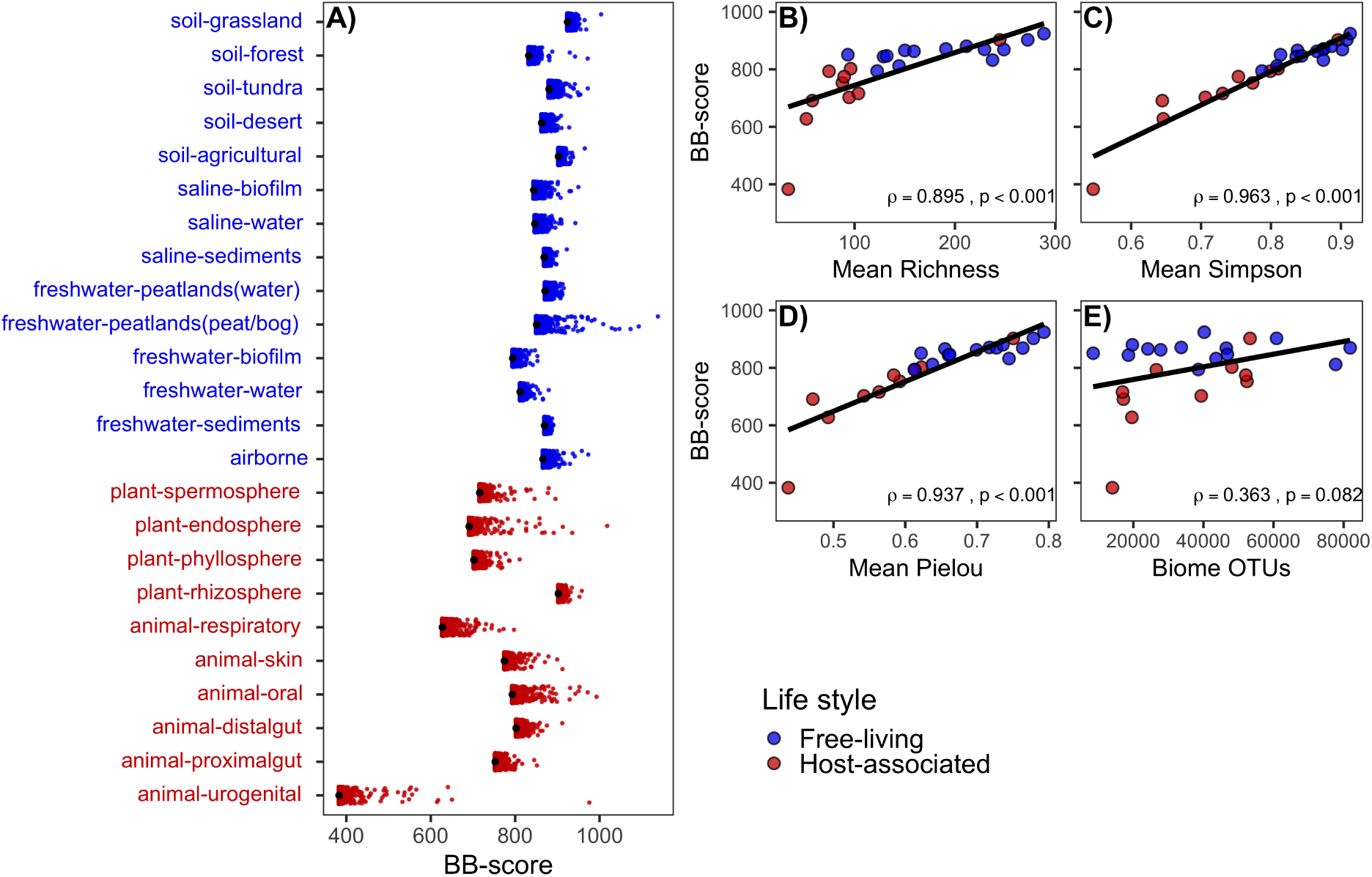
Pooled Baas–Becking score (BB-score) of bacterial species across biomes and its relationship with diversity metrics. (A) Mean BB-scores per species across soil, freshwater, saline, plant-associated, and animal-associated biomes (blue, freeliving; red, host-associated). Black symbols: mean BB-score per biome. (B–E) Relationship between mean biome BB-scores and α-diversity metrics: (B) species richness, (C) Shannon index, (D) Pielou’s evenness, and (E) Simpson index and biome species pool. (F) No significant correlation was found between mean biome BB-score and total number of OTUs per biome. Inset values: Spearman correlation coefficient (ρ) and corresponding p-value.

### Ubiquitous top-performing microbes

We examined OTUs that ranked within the top 5% of BB-scores in individual biomes to distinguish between biome specialists and taxa with high inter-biome performance (Fig. 4). The number of biomes where OTUs were in the top 5% of BB-scores were positively correlated with their mean BB scores across the other biomes (Spearman ρ = 0.611, P < 0.001; Fig. 4A). Most OTUs were among the top performers only in a single or a few biomes, and were restricted in their occurrence to <20 biomes (Fig. 4B).

**Figure 4.**
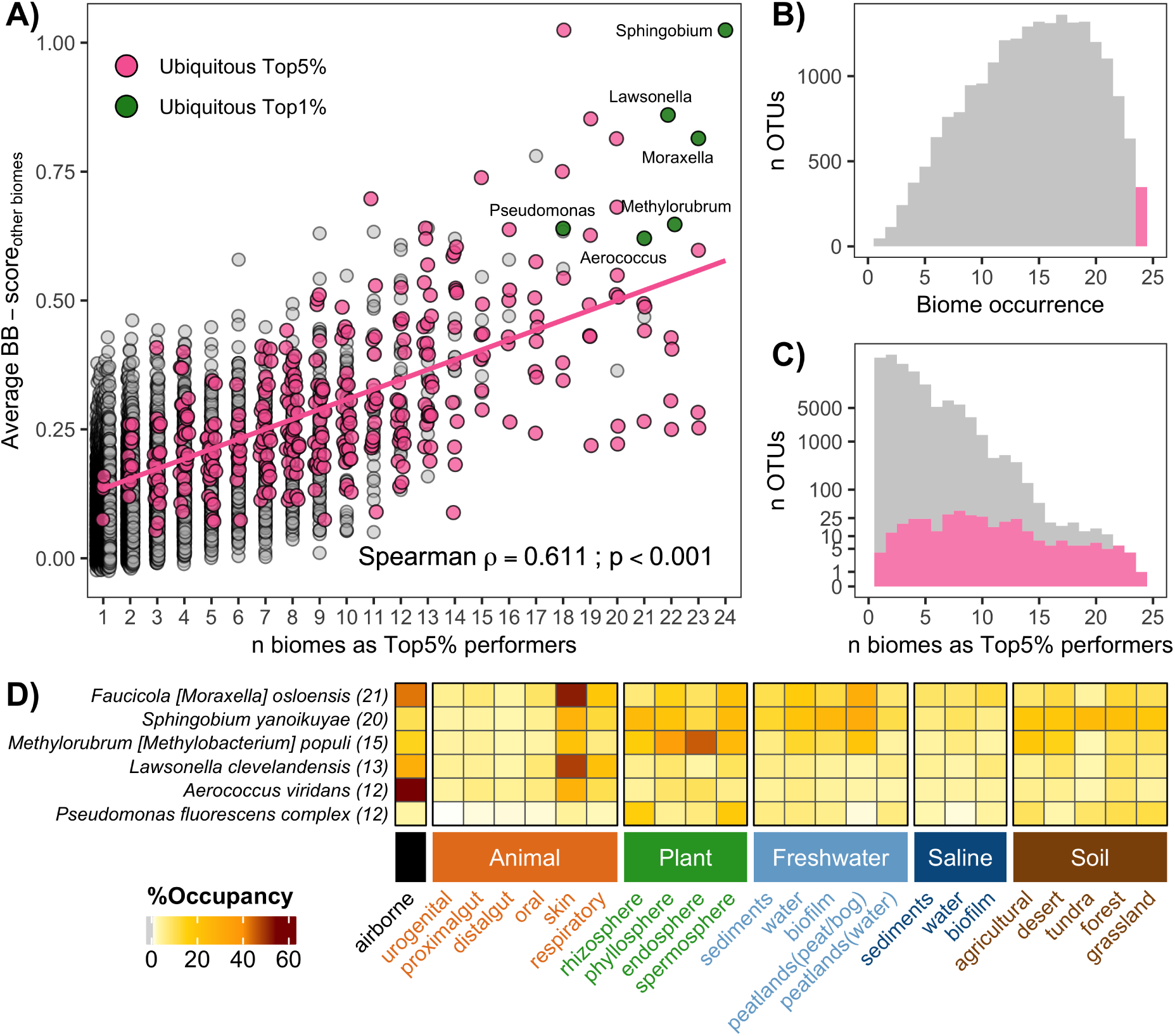
Restricted and ubiquitous top performing bacteria. (A) Inter-biome performance (average normalized BB-score across all biomes) of those OTUs that were in the top 5% of performers in 1 or more biomes. The number of biomes in which OTUs ranked in the top 5% was positively correlated with their inter-biome performance. The subset of ubiquitous microbes detected in all 24 biomes are highlighted in pink. Six ubiquitous microbes were in the top 1% of performers in more than half of the biomes and their average BB-score was >99.9% of all OTUs (highlighted in green). These taxa are annotated as Aerococcus viridans (NCBI accession number: HM813593), Lawsonella clevelandensis (GQ050671), Methylorubrum (previously Methylobacterium) populi (JQ660315; previously Methylobacterium populi), Faucicola (previously Moraxella) osloensis (HM287028), Pseudomonas fluorescens complex (EU449578), Sphingobium yanoikuyae (JF459964). (B) Occurrence of OTUs across biomes that were in the top 5% of performers in 1 or more biomes. Ubiquitous OTUs highlighted in pink. (C) Distribution of the same set of OTUs according to the number of biomes in which they ranked among the top 5% of performers. Ubiquitous winners highlighted in pink. (D) Heatmap of biome-level occupancy (% of samples per biome) of the six top-performing microbes. Numbers in parentheses indicate the number of biomes in which each lineage was ranked within the top 1% of performers. The color bar indicates relative occupancy.

A small set of OTUs consistently ranking among the highest performers were ubiquitously present in all 24 biomes (Figs. 4B, 4C). By further restricting our criteria to ubiquitous OTUs that were amongst the top 1% BB-score performers in 50% or more of biomes, six taxa stood out: *Aerococcus viridans*, *Faucicola* (previously: *Moraxella) osloensis*, *Lawsonella clevelandensis*, *Methylorubrum populi*, *Sphingobium yanoikuyae*, and the *Pseudomonas fluorescens* species complex. Albeit highly successful in 12 to 21 biomes (Fig. 4D), these taxa nevertheless displayed contrasting inter-biome distribution patterns: *M. osloensis* and *L. clevelandensis* had the highest occupancies in animal skin and respiratory system biomes, *P. fluorescens, M. populi* and *S. yanoikuyae* were most frequently found in samples from plant-related biomes, and *S. yanoikuyae* and *P. fluorescens* were also common members of soil and freshwater communities (Fig. 4D, Suppl. Table S3). Interestingly, all these taxa but *P. fluorescens* were also within the top 1% performers of the airborne biome, with the highest occupancies by *A. viridans* (56% of samples), *F. osloensis* (43%), and *L. clevelandensis* (28%).

A bibliometric survey (Web of Science^TM^; Suppl. Table S4) shows that literature coverage is highly uneven among the top-performing taxa: *P. fluorescens* is referenced in >24,000 records, whereas each of the remaining top performers is represented by only a few dozen to several hundred records. By comparison, common pathogens such as *Streptococcus pneumoniae* and *Staphylococcus aureus* are associated with >69,000 and >340,000 records, respectively (Suppl. Table S4).

### Cross-biome connectivity by common top-performing microbes

We constructed a network based on the inter-biome Jaccard similarity of OTUs in the top 5% BB-scores within each biome (Fig. 5A). The resulting network was highly clustered, while average path lengths were comparable to those of randomized networks, yielding a small-worldness index of 1.19 (Suppl. Table S5). Biomes organized into coherent clusters by lifestyle and habitat type. Animal-associated biomes formed a dense module, with proximal and distal gut communities showing strong overlap and close connections to oral, respiratory, skin and urogenital microbiomes. Plant-associated biomes (phyllosphere, rhizosphere, endosphere, spermosphere) also clustered together and exhibited strong connectivity with soils. Freshwater biomes were tightly linked to plant and soil systems but exhibited weaker overlap with top-performing community members in saline and animal-associated biomes. Airborne microbiomes displayed broad overlap with phyllosphere, soil, freshwater and animal biomes, consistent with extensive sharing of top-performing taxa across biome groups.

**Figure 5.**
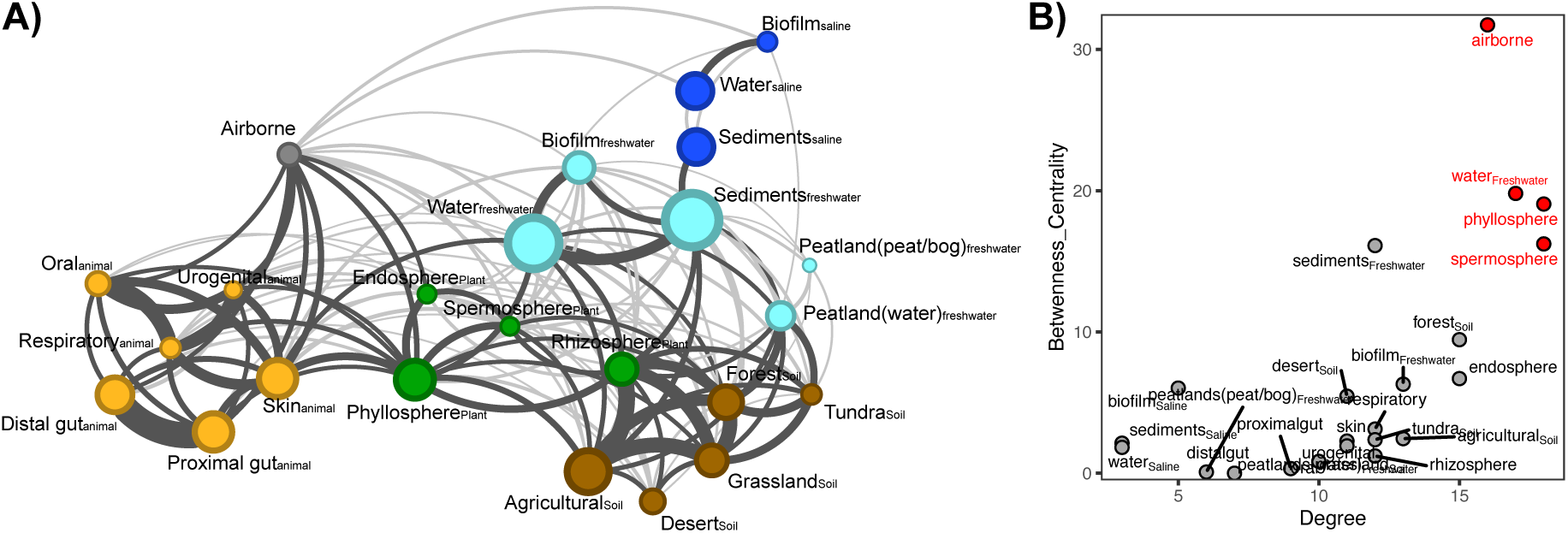
Presence-absence network of biomes based on top-performing OTUs. A) Network constructed from the top 5% of OTUs ranked by BB-score within each biome, using the Jaccard similarity index to quantify shared OTUs. Edges with Jaccard similarity ≥ 0.1 are shown in darkgrey, and those < 0.1 in lightgrey. Edge thickness represents similarity strength (Jaccard similarity), while node size corresponds to the number of top OTUs per biome. To filter edges, we applied the α (co-occurrence) index from Mainali et al. 2022, which estimates the maximum-likelihood deviation (α) from random species co-occurrence; only biome links with α > 0 (more similar than expected by chance) and p-value < 0.05 were retained. Node colors indicate biome type: orange = animal-associated, green = plant-associated, brown = soil-associated, light blue = freshwater-associated, dark blue = saline-associated, black = airborne. B) Relationship between node degree and betweenness centrality. Red points indicate biomes in the upper-right quadrant, identified as keystone biomes representing the major cross-biome connectors.

Among all biomes, the phyllosphere showed the highest network degree, sharing more than 150 top-performing taxa with other biome groups (Suppl. Table S6). In contrast, saline biomes (water and biofilm) displayed the lowest overlap with the remainder of the network. Degree-betweenness centrality analyses further highlighted airborne, freshwater, phyllosphere and spermosphere biomes function as key global connectors for top performers (high betweenness), whereas saline biomes, peatlands and animal gut biomes occupied more peripheral positions with comparatively weak connectivity (Fig. 5B).

## Discussion

Across ∼0.6 million microbial communities spanning 24 biomes, our analyses reveal three main patterns. First, a majority of OTUs cross a fundamental lifestyle boundary, indicating pervasive ecological flexibility at the phylotype level. Second, occupancy and performance are frequently decoupled; many widespread taxa are only moderate performers, whereas a small subset of “true generalists” repeatedly ranks among the top performers across biomes. Third, a network of shared top performers identifies a small set of hub biomes that disproportionately structure cross-biome connectivity, with salinity acting as a major barrier.

Global analysis revealed that about 64% of microbes cross the boundaries between host-association and the free-living lifestyle and that such ecological flexibility is the rule rather than the exception (Fig. 2A). This pattern is consistent with frequent transitions and/or repeated dispersal between host-associated and free-living contexts, highlighting the role of multicellular eukaryotes as transient habitats or dispersal vectors (23) that increase biome connectivity. For example, members of the genus *Limnohabitans*, widespread free-living freshwater bacteria, can also transiently occur in the gut of zooplankton (24). Consequently, host-associated microbes appear far more dispersed and diverse than previously recognized, challenging observations that host-associated assemblages are typically less diverse than their free-living counterparts (4).

Occurrence in single biomes was rare (Fig. 2B). The low proportion of apparent specialists for the airborne biome (0.27%) likely represent the lower limits of quantification within our dataset (Suppl. Table S2); exclusive specialization to this biome is not plausible, and the scarcity of “air-only” OTUs rather reflects the integrative nature of the atmosphere which samples microbes from multiple sources. The transient nature of seeds as habitats that only persist until germination (25) and the acidic, anoxic, and nutrient-poor conditions of peatlands (26) possibly preclude the establishment of biome-specialized microbes. By contrast, the greatest biome-specialist occurred in saline biomes (aquatic and sediments combined with ∼ 7%). Salinity gradients between these and other habitats, likely favor the selection of specialized lineages and higher fractions of biome-unique microbial diversity(27).

By ranking OTUs against their local competitors across communities, BB-scores yield a context-dependent measure of relative performance that complements prevalence-only or abundance-only summaries that only capture one dimension of ecological success. As a relative metric defined on specified metacommunities (here: biomes), the BB-score enables OTU-specific comparisons across space (and, where available, time) in a community context. Tight correlations between biome-mean BB-scores and classical alpha-diversity metrics (Fig. 3B–E) indicate that the metric is consistent with established community-level descriptors while adding a taxon-centered axis of inference.

Biome occurrence and ecological success (BB-score) were decoupled: widely distributed OTUs were often moderately ranked, and most taxa were top performers in only few biomes (Fig. 4B, C), indicating that broad occupancy reflects persistence rather than disproportionally high competitive success. The large number of restricted winners, compared to widely distributed ones, suggests that performance is likely shaped by trade-offs between adaptation and growth capabilities, consistent with specialist–generalist reasoning (28). Yet a small subset of lineages increasingly ranked among the top across multiple biomes while sustaining high inter-biome performance (Fig. 4A), indicating the presence of ubiquitously successful taxa that might be termed “true generalists”.

Such truly generalist microbes were rare, and only few bacterial lineages ranked within the top 5% of BB-scores across all biomes (Fig. 4). Among these, several are opportunistic human pathogens (*A. viridans, F. osloensis, L. clevelandensis*), while others are frequently isolated from environmental samples such as plants or soils (*M. populi, S. yanoikuyae*, *P. fluorescens* complex). Despite their ecological success, most globally top-performing taxa are poorly studied compared to the “classic” human pathogens, highlighting our anthropocentric focus on microbiology, with clinical threat by far outweighing environmental success (Suppl. Table S4). Our shortlist of ubiquitously successful microbial species with high inter-biome transmissibility (*bona fide* “Baas-Becking’s demons”) may thus represent promising microbial models for One Health and biotechnological research (29).

To explore genomic correlates of broad ecological success, we performed a targeted pangenome comparison for a small set of BB-score winners that could be unambiguously linked to GTDB species with sufficient genome representation. In this limited analysis, the two globally top-performing taxa exhibited larger rarefied pangenomes than three biome-restricted winners, suggesting a possible association between cross-biome success and expanded accessory gene repertoires (Suppl. Fig. S1).

Network analysis of shared top performers revealed a small-world pattern, with biomes clustering into coherent modules globally connected by a few hub biomes: phyllosphere, atmosphere, and freshwater (Fig. 5). Living plants account for 80% of global biomass (30) and their leaf surface area is approximately double the land surface area (31), representing one of the planet’s largest microbial interfaces. Plant leaf surfaces constitute extensive yet selective habitats (32) that can facilitate microbial exchange among animals, soils, and freshwater systems.

The atmosphere functions as a dispersal corridor, linking successful microbes associated with animals and plants rather than acting as a reservoir. Virtually all airborne OTUs also occurred in other biomes (Fig. 2B), consistent with prior findings that the aerobiome is dominated by microbes sourced from other environments (33, 34). The high occupancy of top-performing bacteria in airborne samples (Fig. 4D) and the high betweenness centrality of the airborne biome (Fig. 5B) both support its role as a bridge linking microbiologically separated biome clusters. Freshwater ecosystems are often interconnected by natural or artificial waterways that transport organisms across spatial gradients and link otherwise isolated habitats within landscapes and regions (35, 36). Terrestrial-aquatic interfaces, moreover, facilitate dispersal, as soil and stream microbial communities are often coupled via hydrological transport (37). The high representation of global microbial diversity in freshwater biomes (∼60%; Suppl. Fig. S2D) and high betweenness centrality of top-performing lacustrine taxa (Fig. 5B) together highlight their role as reservoirs of broadly adapted microbial lineages. Beyond simple linkage, the hydrological networks and directional flow can repeatedly mix source communities and export taxa downstream, generating persistent corridors that connect soils, vegetation, inland waters, and (via downstream transport) coastal systems.

Limited overlap of top-performing taxa between saline and freshwater biomes and the highest proportions of biome-specific OTUs (Fig. 2B) position salinity as a major biogeographic barrier shaping microbial distributions (Fig. 5A). These patterns reflect differential microbial success across the freshwater-saline boundary (27, 38). However, saline sediments shared a substantial fraction of top-performing microbes with their freshwater counterparts (Fig. 5A). This agrees with observations that sediment microbiota in Lake Baikal had high compositional similarity to marine benthic assemblages (39). Constraints other than salinity, such as organic carbon availability and quality, or redox gradients, may act as dominant environmental filters in such environments (40).

Our study was based on phylotypes defined as 16S rRNA OTUs at 99% identity (21), which approximate species-like units but do not resolve strain-level variation (41). In addition, reference and sampling biases (notably toward animal-associated microbiomes; Fig. 1), cross-study heterogeneity in protocols, and detection limits - particularly in low-biomass samples - can influence apparent occupancy and relative performance despite standardization. Finally, biomes were defined operationally (Fig. 1, Suppl. Table S1) and encompass heterogeneous conditions. However, systematic differences in diversity metrics and in BB-score distributions across biomes (Fig. 3B-C) confirm these shared ecological constraints, supporting biomes as defined as an effective integrative scale for the macroecological analysis of microbes (4, 42, 43). Our analysis might also contribute to a better understanding of the environmental reservoirs of clinical threats, as exemplified by the WHO Bacterial Priority List (44) or the ESKAPE pathogens (45). For example, *Pseudomonas aeruginosa,* known for its occurrence in numerous environments (46), ranked within the top 5% taxa in 14 of the 23 biomes in which it was detected (Suppl. Table S7), including desert soil (47), the phyllosphere (32), or marine biofilms (48). Likewise, *Streptococcus pneumoniae* again occurred in 23 biomes; beyond established host-associated habitats, it also ranked within the top 5% of BB-scores in plant-associated biomes and among lacustrine waterborne bacteria (Suppl. Table S7).

Altogether, we provide a novel macroecological framework for assessing microbial success across biomes. By combining large-scale occupancy patterns with a novel taxon-centered performance index, we show that ecological flexibility is widespread, that strictly biome-restricted OTUs constitute a small fraction of local diversity, and that a few taxa repeatedly rank among top performers across multiple environments. By identifying both high-performing taxa and the hub environments and transport processes that couple biomes, our results outline selective pathways by which dispersal and environmental filtering jointly shape global microbiome structure. As genomic resources expand, integrating the BB-score with genome-resolved traits will help advance toward a more unified and mechanistic framework for microbial macroecology.

## METHODOLOGY

### Data collection

We used the Microbe Atlas Project (MAP) dataset that contains operational taxonomic units (OTUs) of microbial 16S rRNA genes from approximately 2 million samples generated by amplicon or metagenomic sequencing (21). OTUs were defined at an identity threshold of 99% (approximating “species” level resolution) using MAPseq (22). Samples with (i) a minimum of 1000 quality-filtered reads, (ii) at least 3 different OTUs, and (iii) valid biome descriptors in the MAP metadata were further retained for downstream analyses (Suppl. Table S1). A dataset of 576,531 communities was maintained after filtration and classification, containing 124,772 distinct OTUs (Suppl. Table S1). Extremely rare OTUs with occurrences <0.01% in single biomes were discarded from downstream analyses.

We split biomes into two sets of distinct microbial lifestyles with respect to macroecological rules (49) and community assembly processes (50), i.e., free-living and host-associated. Free-living biomes comprise habitats in which species grow independently of single multicellular hosts, such as various soil biomes, lacustrine and marine waters, sediments, biofilms, peatlands, and the airborne microbiome. Host-associated biomes were defined as environments where organisms reside on or within animals or plants and experience strong biotic filters, such as host-derived nutrients and physical niches (51) and included animal epithelia and digestive tract (urogenital, oral, skin, respiratory, proximal gut, distal gut) as well as plant surfaces, tissues, and seeds (rhizosphere, phyllosphere, endosphere, spermosphere).

### Occurrence across biomes

To compare OTU occurrence (presence/absence) across biomes, we first standardized our sampling to the biome with the lowest sample number (plant spermosphere, n=1059). We then randomly selected 1059 samples (without replacement) from each of the 24 biomes and recorded the biomes in which each OTU was detected. This subsampling procedure was repeated 1,000 times. Finally, for each OTU, the total (cumulative) number of distinct biomes in which it ever appeared was recorded. “Single-biome OTUs” were defined as those OTUs that were exclusively found -once or multiple times-in the same single biome over all iterations.

### Elo-rating and Baas-Becking (BB)-score

We quantified microbial species relative performance in each of the 24 custom-defined MAP biomes using a modified version of Elo-rating for multiplayer ranking, as implemented in multielo v0.4.0 (https://github.com/djcunningham0/multielo). We first applied the original multiplayer Elo-rating as used in experimental microbial metacommunities (19). However, classic Elo-rating maintains zero-sum properties, and “players” not participating in “matches” (i.e., taxa absent from communities) do not change rank (Suppl. Fig. S3, Suppl. Table S8). This limitation reduces ecological realism because species absence in a community implies both abiotic and biotic filters affecting species colonization and growth, therefore their relative performance.

Our proposed variation of rating score considers the relative abundance (performance) of all OTUs present in a given sample for updating their rating, but at the same time introduces a penalization of OTUs detected in the biome pool (or “tournament”) but absent from the focal community (“match”). Specifically, each absent OTU was assigned the rank of the lowest-performing OTU in that sample, and its overall rank was updated accordingly. This absence-correction breaks the strict zero-sum property of classical Elo-rating. Since this way of calculating scores assumes that every species occurring in a biome could potentially be present in every sample, “but, the environment selects” (20), we named our absence-corrected variant of the Elo-rating the “Baas-Becking-score” (BB-score).

Both Elo-rating and BB-score use the same scoring function (*S_observed_*), defined by a decay function fitted to the distribution of OTUs in each biome (equation 1):

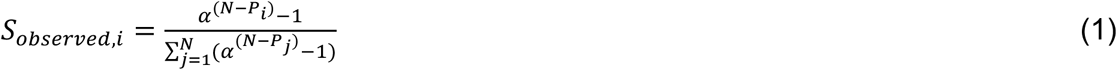

where *N* is the number of total OTUs in the biome, *P_i_* is the rank position of OTU *i*, *α* is the base of the scoring function, and *j* indexes all OTUs in the biome. The parameter *α* was estimated independently in each biome using the function “nlsLM” from the R-package minpack.lm (52). For inter-biome comparison, the average *α* of all biomes, corresponding to 1.017, was used (Suppl. Table S2).

For rating calculations, each sample was operationally considered as a “match” among all OTUs detected in that sample. For BB-scores, absent OTUs were penalized as outlined above. All OTU ratings were updated after each match according to equation 2:

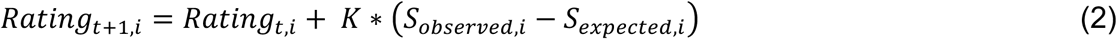

where K is a gain parameter (18, 53) (a constant, K=10), S_observed,i_ is the observed score assigned to OTU_i_ in that match (according to equation 1), and S_expected,i_ is the expected score given the current ratings of all participants divided by the total number of pairwise interactions following equation 3:

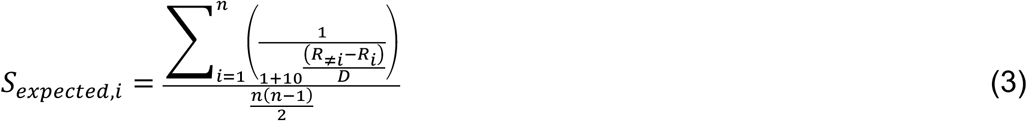

where *D* is a constant (54) (D=400), *n* is the number of OTUs in that match, *R_i_* is the current rating of the *OTU i* and *R*_≠*i*_ is the current rating of each OTU competing against *OTU_i_*.

For Elo-rating and BB-score calculations, all OTUs within a biome were initialized at a rating of 1,000. Contrary to the Elo-rating, which follows zero-sum properties, the BB-score estimations are influenced by the number of matches (samples per biome; as exemplified in Suppl. Fig. S4), therefore, each biome data set was randomly subsampled to the smallest biome size (1059 samples, without resampling; Suppl. Table S2). This process was repeated 1,000 times and the average rating per OTU was used for downstream analysis. Calculations were computed with a custom Python script, MultiElo_switchable.py, available at https://github.com/angelrainf/microbes.bbscore. The script was run using the --mode “classic” and “corrected” for Elo-rating and the BB-score estimation, respectively.

### Average biome performance of top-performing OTUs

To identify disproportionally successful taxa, we focused on OTUs in the top 5% of BB-scores in one or several biomes. For the quantification of the performance of these OTUs across the other biomes, we calculated average BB-scores for all biomes where they did not fall within the top 5% of BB-scores as follows:

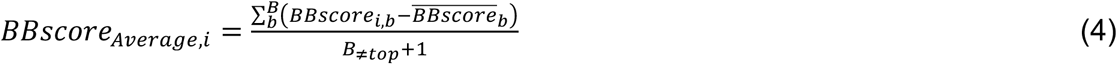

where *BB_scorei,b_* is the score of the OTU *i* in the biome b, 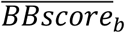 is the mean BBscore of all OTUs in the biome *b, and B*_≠*top*_ is the total number of biomes where the OTU *i* is not in the top5%. Thus, high average BB-score values indicate that OTUs consistently ranked highly across biomes where they were not in the top 5%.

### Statistical analysis

All statistical analyses were conducted in R v4.1.2 (55). Visualization used package ggplot2 (version 3.5.1). Richness and Simpson (1-D) diversity were computed using vegan (56). Pielou’s evenness was calculated by dividing the Shannon index by the natural log-transformed richness. Associations between biome-level BB-scores and diversity indices (richness, Simpson, Pielou’s evenness) were tested with Spearman’s rank correlation and differences between free-living vs host-associated biomes average BB-scores were assessed with a Wilcoxon rank-sum test.

The biome connectivity network used the presence/absence of the top 5% BB-scores OTUs within each biome using the Jaccard dissimilarity. To determine whether similarity was higher than by chance, we used the maximum likelihood estimation, α, from the CooccurrenceAffinity R package (57). P-values from the dissimilarity tests were corrected for multiple comparisons using the Bonferroni procedure. Edge weights were defined as Jaccard similarity (1 - Jaccard dissimilarity). Pairs were filtered to those with α > 0 (indicating higher similarity than expected by chance), and p-adjusted < 0.05. Transitivity, average path length and small-worldness were calculated using qgraph package (58). Small-worldness index significance was assessed by a Monte Carlo randomization test (59) against 1000 degree-preserving null networks. The final network was visualized in Gephi.

## Acknowledgments

We thank Janko Tackmann for assistance with the Microbe Atlas Project dataset. This study was funded by the Swiss National Science Foundation Grant Number 10000877.

## Data availability

Code to reproduce the results and Python functions for rating estimations are available in the online repository https://github.com/angelrainf/microbes.bbscore. Datasets, including rating and occurrence estimates, are available via Zenodo (doi:10.5281/zenodo.18432582).

