## Supplementary Material for "Macroecology of microbial performance across Earth’s biomes"

**This PDF file includes:**

Supplementary Material

Supplementary Figures S1 to S4

**Supplementary Material**

**Targeted genome selection and pangenome analysis of BB-score winners**

To provide an exploratory, genome-resolved complement to BB-score–based performance patterns, we assessed whether a subset of globally successful (“true generalist”) taxa shows evidence for larger pangenomes compared with biome-specific winners. Because the BB-score is computed on 16S rRNA OTUs, this analysis was limited to taxa for which OTU-to-genome mapping was sufficiently unambiguous and for which adequate genome representation existed for pangenome estimation. We defined two categories of taxa based on BB-score patterns:

1. Globally successful taxa (“true generalists”) are identified as top-performing across multiple biomes and exhibiting high inter-biome performance.
2. Biome-specific winners are defined as taxa with strong performance concentrated within particular biomes.

To connect OTU-based taxa to genome-defined species, we retained only cases with a clear association to a GTDB species (i.e., avoiding ambiguous species-complex assignments). We further required at least eight genomes per species to enable rarefaction-based pangenome estimation. For globally successful taxa, we additionally required that genomes were derived from more than one biome/origin category based on available metadata, to reduce reliance on single-context genome collections.

This filtering yielded two globally successful species and three biome-specific species:

1. Global winners (true generalists): *Sphingobium yanoikuyae* and *Methylorubrum populi*.
2. Biome-specific winners: *Treponema_B pedis*, *Alteromonas stellipolaris*, and *Erwinia mallotivora*.

Several additional top-performing taxa could not be included due to either (i) ambiguous mapping to GTDB species complexes (e.g., *Sinorhizobium meliloti*, *Pseudomonas fluorescens* complex), or (ii) insufficient genome counts (n < 8 genomes; e.g., *Entomoplasma ellychniae*, *Aerococcus viridans*). In one case (*Lawsonella clevelandensis_A*), the number of genomes was sufficient, but all genomes originated from host-associated environments, and the taxon was therefore not used as a comparator for globally distributed taxa under our biome-diversity criterion.

**Pangenome analysis**

Genome accession identifiers were obtained from GTDB species entries. Corresponding nucleotide and annotation files were downloaded from NCBI. Protein-coding genes were predicted using Prodigal v2.6.3 (1) with default parameters. Pangenomes were computed using PPanGGOLiN v2.2.6 (2) (ppanggolin all) using the built-in rarefaction option to estimate how pangenome size (gene families) accumulates as a function of the number of genomes sampled per species. Rarefaction outputs were exported as rarefaction.csv and processed in R v4.1.2 (3) using custom scripts to compute the median and interquartile range (IQR) of pangenome size across rarefaction replicates at each sampling depth (genomes_count).

**Supplementary Results**

**Exploratory pangenome comparison of global and biome-specific BB-score winners**

After applying the OTU-to-GTDB mapping and genome-availability filters (Supplementary Methods), we retained two globally successful taxa with sufficient genome representation and non-single-biome genome collections (Genome accessions are listed in Supplementary Table S9.):

- *Sphingobium yanoikuyae* (27 genomes; median genome completeness 100%; median GC 65%; median coding density 90%).
- *Methylorubrum populi* (10 genomes; median completeness 100%; median GC 69%; median coding density 86.3%).

For biome-specific winners, three species met the minimum genome requirement:

- *Treponema_B pedis* (9 genomes; median completeness 99.78%; median coding density 88.5%; median GC 37%).
- *Alteromonas stellipolaris* (19 genomes; median completeness 100%; median coding density 88.6%; median GC 44%).
- *Erwinia mallotivora* (8 genomes; median completeness 100%; median coding density 86.2%; median GC 52%).

**Pangenome rarefaction patterns**

PPanGGOLiN rarefaction analyses showed that pangenome size increased with the number of genomes sampled for all five taxa considered (Suppl. Fig. S1). Because genome availability differed among species (8–27 genomes), we compared pangenome sizes under a standardized sampling depth by truncating rarefaction curves at eight genomes, corresponding to the maximum available for the smallest genome set (*Erwinia mallotivora*). Under this matched sampling depth, the two globally successful taxa (*Sphingobium yanoikuyae* and *Methylorubrum populi*) exhibited substantially larger pangenomes than the three biome-specific winners (*Treponema_B pedis*, *Alteromonas stellipolaris*, *E. mallotivora*) (Suppl. Fig. S1a). Specifically, when sampling eight genomes per species, estimated pangenome sizes were 12,706 gene families for *S. yanoikuyae* and 11,796 for *M. populi*, compared with 6,155 (*E. mallotivora*), 5,938 (*A. stellipolaris*), and 5,655 (*T. pedis*) gene families for the biome-specific winners. Thus, in this limited comparison, rarefied pangenome sizes of the global winners were approximately two-fold larger than those of the biome-specific winners.

Across the full available genome sets, pangenome size continued to increase with additional genomes, but the magnitude of this increase differed among taxa (Suppl. Fig. S1). Notably, *S. yanoikuyae* showed sustained pangenome expansion as sampling increased from two genomes to the full set of 27 genomes, reaching an estimated pangenome size of 25,613 gene families at 27 genomes. In contrast, for the three biome-specific winners, pangenome sizes remained below approximately 8,000 gene families even when using their full genome sets (8–19 genomes).

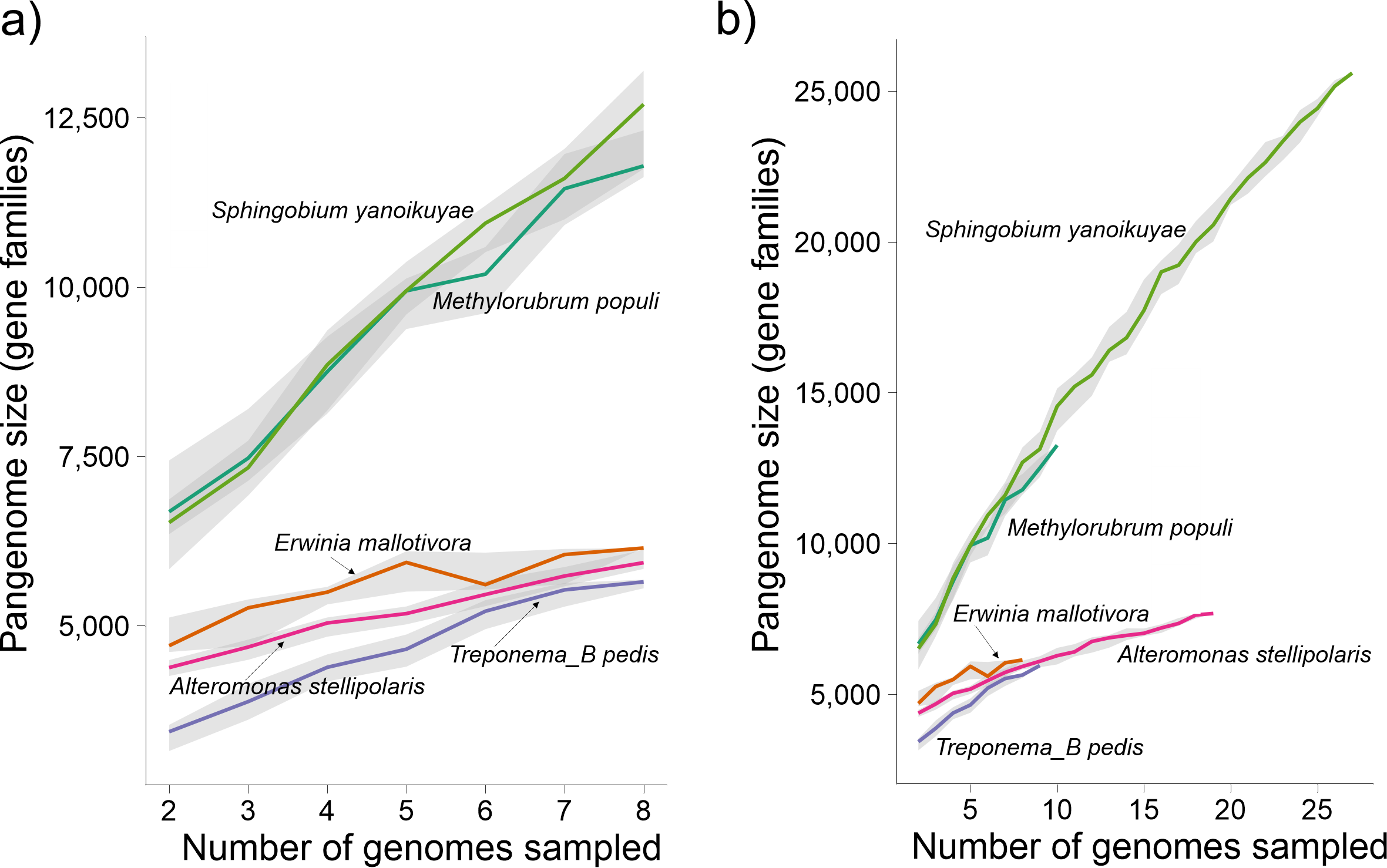

**Supplementary Figure S1.** Pangenome rarefaction of globally successful and biome-specific BB-score winners. Pangenome size, quantified as the number of gene families, is plotted against the number of genomes sampled per species using PPanGGOLiN rarefaction. Colored lines show the median pangenome size across rarefaction replicates, and the grey area represent the interquartile range. The two globally successful taxa (*Sphingobium yanoikuyae* and *Methylorubrum populi*) are compared with three biome-specific winners (*Erwinia mallotivora*, *Alteromonas stellipolaris*, and *Treponema_B pedis*). (a) Curves truncated to eight genomes per species for direct comparison under matched sampling depth (eight genomes correspond to the maximum available for the smallest genome set). (b) Curves are shown for the full number of genomes available for each species to illustrate continued pangenome expansion with additional genomes.

**Supplementary Discussion**

**Larger accessory gene repertoires may facilitate cross-biome success**

This targeted genome-resolved comparison provides suggestive evidence that globally top-performing BB-score taxa may harbor larger pangenomes than biome-restricted winners. Larger pangenomes and stronger pangenome expansion under rarefaction are consistent with the idea that broad ecological success can be supported by expanded accessory gene repertoires, potentially increasing metabolic versatility, stress tolerance, and the ability to exploit heterogeneous resources across environments (4).

At the same time, this analysis is intentionally hypothesis-generating and should be interpreted cautiously. First, the comparison includes only two global winners and three biome-restricted winners, limiting generality and precluding strong statistical inference. Second, pangenome size is sensitive to genome sampling and to the diversity of isolates available in public repositories; although we used the lowest genome number (n = 8) to improve comparability, residual biases may remain. Third, OTU-to-species mapping is not always one-to-one (particularly for species complexes), restricting the set of taxa that can currently be linked confidently from amplicon-defined OTUs to genome-defined species.

Future work that scales genome-resolved analyses to a larger set of BB-score winners, including systematic controls for phylogeny and genome size and incorporation of functionally annotated gene categories, will be required to evaluate whether expanded pangenomes are a consistent feature of “true generalists” and to identify which functional capacities most strongly predict cross-biome performance.

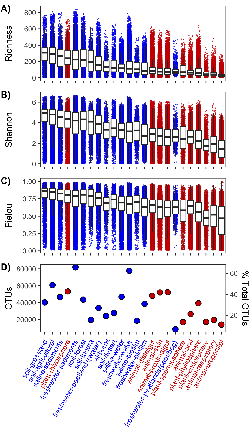

**Supplementary Figure S2. Alpha- and gamma-diversity across biomes.** (A) Sample-level richness (OTU count), (B) Shannon diversity, and (C) Pielou’s evenness by biome. Boxplots show the median (center line) and the first and third quartiles (bottom and upper limits of boxes); whiskers span the 10^th^ and 90^th^ percentiles. Overlaid points represent the individual samples. (D) (gamma)γ-diversity (total OTUs observed across all samples within each biome); each point represents one biome. Biomes on the x-axis are ordered by the median richness in (A). Colors denote lifestyle classification: blue, free-living biomes; red, host-associated biomes.

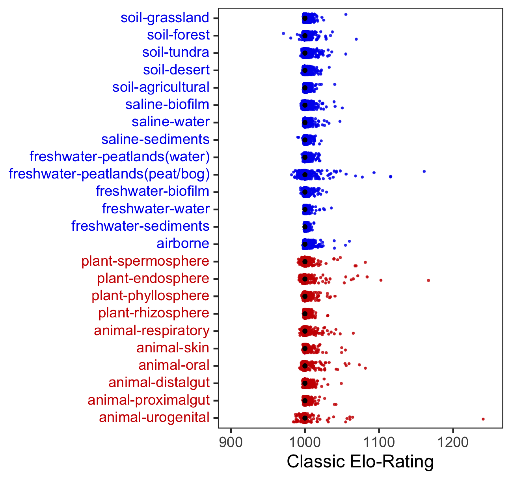

**Supplementary Figure S3. Elo-ratings across biomes.** Each dot represents the Elo-rating of an individual OTU across soil, freshwater, saline, plant-associated, and animal-associated biomes (blue: free-living; red: host-associated). Black dots indicate the average Elo-ratings per biome, which converges at 1000 due to the zero-sum game property of Elo-ratings estimations, where per community gains by some OTUs are balanced by losses of others.

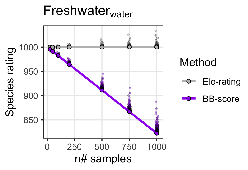

**Supplementary Figure S4. Sample-size dependence of species rating estimations.** Comparison of Elo-rating and BB-score estimates using freshwater–water biome data with increasing sample sizes (n = 25, 50, 100, 200, 500, 750, 1000). Average Elo-ratings remain stable regardless of sample number, whereas BB-scores decline systematically with sampling depth due to penalization of absences.
